# Inhibition of Wnt signalling by Notch via two distinct mechanisms

**DOI:** 10.1101/2020.04.14.037788

**Authors:** Ahmet Acar, Ana Hidalgo-Sastre, Michael K. Leverentz, Christopher G. Mills, Simon Woodcock, Martin Baron, Giovanna M. Collu, Keith Brennan

## Abstract

Notch and Wnt are two essential signalling pathways that help to shape animals during development and to sustain adult tissue homeostasis. Although, they are often active at the same time within a tissue, they typically have opposing effects on cell fate decisions. In fact, crosstalk between the two pathways is important in generating the great diversity of cell types that we find in metazoans. However, several different mechanisms have been proposed that allow Notch to limit Wnt signalling, driving a Notch-ON/Wnt-OFF state. Here we explore these different mechanisms in vertebrate cells and demonstrate two distinct mechanisms by which Notch itself, can limit the transcriptional activity of β-catenin. At the membrane, independently of DSL ligands, Notch1 can antagonise β-catenin activity through an endocytosis mediated mechanism that is dependent upon its interaction with Deltex and sequesters β-catenin into the membrane fraction. Within the nucleus, the intracellular domain of Notch1 can also limit β-catenin induced transcription through the formation of a complex that requires its interaction with RBPjκ. We believe these mechanisms contribute to the robustness of cell-fate decisions by sharpening the distinction between opposing Notch/Wnt responses.

## Introduction

Animal development and adult tissue renewal are controlled by a handful of different signalling pathways that define a bewildering array of different cell types (Pires-daSilva and Sommer, 2003). This clearly raises the question of how so few signalling pathways can generate such diversity. The answer lies, in part, in crosstalk between the signalling pathways that allows one pathway to regulate or alter the output of another. The Notch and Wnt signalling pathways are a good example of this (Muñoz-Descalzo and Martinez Arias, 2012). These pathways are often active at the same time to regulate the development and maintenance of a particular tissue but frequently have opposing effects on cell fate specification (Hayward et al., 2008). Therefore, it is not surprising that signalling through the two pathways is under strict control to prevent a conflict between them and to resolve into either Wnt-ON/Notch-OFF or Notch-ON/Wnt-OFF. This is particular, important to allow the progression of a cell along particular lineage where subsequent steps are controlled by Wnt and Notch signalling. For example, Notch signalling promotes the differentiation of stem cells in both the skin and the mammary by inhibiting the Wnt signal that supports the self-renewal of these stem cells and will promote the differentiation process (Zhu and Watt, 1999; Lowell et al., 2000; Bouras et al., 2008; Zeng and Nusse, 2010). Similarly, resolving competition between the two pathways is required for the successful differentiation of enterocytes and secretory cells within the intestinal epithelium (Fre et al., 2005; Stanger et al., 2005; van Es et al., 2005b; van Es et al., 2005a).

It is well established that the key parameter of the Wnt signalling pathway is the stability and localisation of the soluble pool of β-catenin (Gottardi and Gumbiner, 2001; Logan and Nusse, 2004; Reya and Clevers, 2005; MacDonald et al., 2009). In the absence of Wnt ligand, the free cytosolic β-catenin interacts with a destruction complex, comprising CK1, APC, Axin and GSK3β, which phosphorylates β-catenin to target it for degradation via the proteosome. The binding of Wnt ligand to the cell surface receptors Frizzled and LRP recruits the cytoplasmic adaptor protein Dishevelled and the destruction complex to the membrane. Consequently β-catenin is no longer phosphorylated and accumulates in the cytosol and nucleus. Within the nucleus, β-catenin induces the expression of downstream target genes by interacting with members of the TCF/LEF family of DNA binding proteins. Notch signalling is triggered by the interaction of Notch receptors and DSL (Delta, Serrate, Lag2) ligands on adjacent cells (Bray, 2006). This leads to the proteolytic cleavage of Notch to release the intracellular domain (NICD), which translocates to the nucleus to induce target gene transcription in a complex with the DNA binding protein RBPj and the transcriptional co-activator Mastermind-like (MAML).

Over recent years, several mechanisms have been proposed to explain the crosstalk between the Notch and Wnt pathways that allow signalling through the two pathways to be resolved into Notch-ON/Wnt-OFF. Studies in Drosophila and mammalian stem and cancer cells have suggested that Notch present within the plasma membrane can modulate the amount and transcriptional activity of Armadillo/β-catenin (Hayward et al., 2005; Sanders et al., 2009; Kwon et al., 2011) by associating with Armadillo/β-catenin present at the membrane and promoting its degradation through endosomal trafficking. In contrast, several other studies in Xenopus, mice and cell lines have suggested that the inhibition of Wnt signalling is mediated by the nuclear form of Notch, NICD, and may require the expression of downstream targets (Nicolas et al., 2003; Hayward et al., 2005; Deregowski et al., 2006; Acosta et al., 2011, Kim et al., 2012). However, it is difficult to know from these works whether one or both proposed mechanisms are required for Notch to affect the crosstalk on β-catenin, as many of the experiments have been completed with forms of the Notch protein that can localise to the membrane upon synthesis and the nucleus after γ-secretase mediated cleavage.

Here, we demonstrate that both a signalling inactive form of Notch that is restricted to the membrane fraction and a nuclear form of Notch, NICD, can inhibit Wnt/β-catenin signalling, although there is a mark difference in their potency. The membrane-restricted form of Notch sequesters β-catenin to the membrane fraction preventing its interaction with TCF/LEF and requires endocytosis to inhibit Wnt signalling. The nuclear form of Notch can also disrupt the interaction between β-catenin and TCF/LEF by forming a complex with β-catenin, but does not require the transcription of downstream Notch target genes. Interestingly, we find that the interaction between β-catenin and nuclear Notch is stabilised by RBPj, although RBPj cannot form a complex with β-catenin itself. This may explain the marked difference in potency between the two different forms of Notch, as RBPj is nuclear restricted. Together, our results indicate that the interaction between Notch and β-catenin can limit Wnt signalling to establish a Notch-ON/Wnt-OFF state to potentially allow robust cell-fate decisions during embryonic development and tissue homeostasis.

## Results

### Notch modulates the transcriptional activity of β-catenin

To determine whether both a membrane localised and a nuclear form of Notch can inhibit Wnt signalling, we initially generated plasmids expressing two distinct Notch proteins, ΔEGF_N1 and NICD. The ΔEGF_N1 construct lacked the 36 extracellular EGF-like repeats, preventing ligand binding, but retained the LNR domain that masks the S2 cleavage site (Fig 1A). As expected, ΔEGF_N1 protein underwent S1 cleavage which occurs during protein maturation (Fig. 1B), was found within the membrane fraction (Fig. 1C), and failed to induce the expression of an RBPj-dependent reporter gene (Fig. 1D). In contrast, the NICD construct, which comprises the intracellular domain only (Fig. 1A), was expressed as expectedly (Fig. 1B), found exclusively within the nucleus (Fig 1C) and robustly activated RBPj-dependent transcription (Fig 1D). Expressing either of these proteins with the Wnt1 protein by co-transfection reduced the response of HEK293T cells to Wnt1 (Fig. 2A); Wnt signalling was monitored in the HEK293T cells by co-transfecting them with the TCFAdTATA reporter plasmid, which contains a minimal adenoviral TATA box regulated by four TCF/LEF binding sites and activate the Wnt pathway more potently than the TOPflash reporter plasmid (Fig. S1A&B). There was, however, a marked difference in the potency of the two Notch proteins with NICD reducing Wnt signalling more markedly. To establish where in the Wnt pathway the two Notch proteins were regulating signalling, we activated the Wnt pathway at different points by expressing S45Fβ-catenin (a stabilised form of β-catenin; (Vecsey-Semjen et al., 2002)) or LEF1-VP16 (a constitutively active form of the LEF1 transcription factor; (Aoki et al., 1999)). ΔEGF_N1 and NICD were both able to inhibit S45Fβ-catenin but neither could reduce LEF1-VP16 induced transcription (Fig. 2B&C) ΔEGF_N1 and NICD similarly regulated the Xenopus β-catenin and Tcf3-VP16 proteins (Fig. S2A&B). These results indicate that both a membrane-restricted and a nuclear form of Notch can reduce Wnt signalling and that it appears to do so at the level of β-catenin.

**Figure 1.**
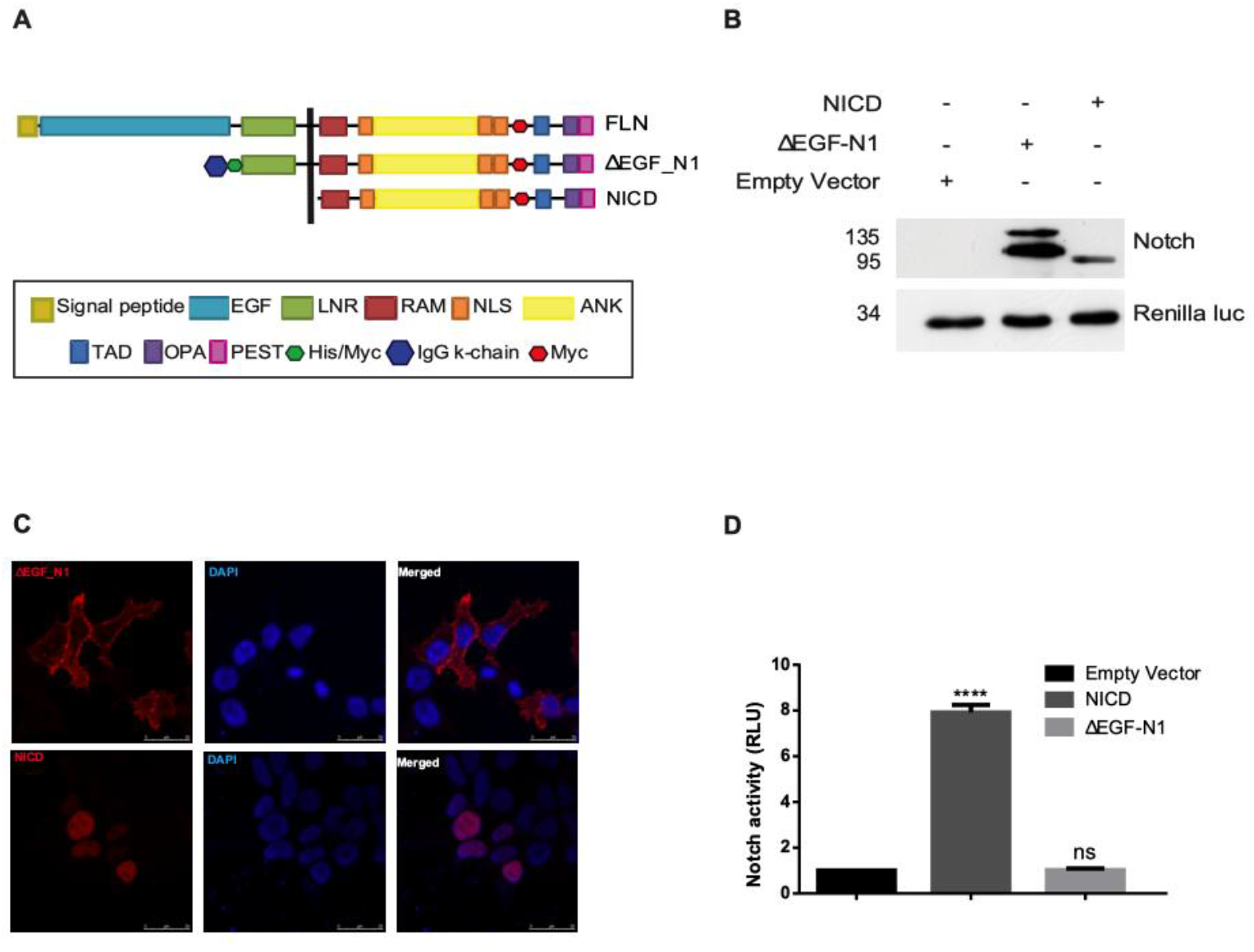
Characterization of ΔEGF_N1 and NICD constructs. **(A)** Schematic showing the structure of the full length, ΔEGF_N1 and NICD proteins. **(B)** Western blot analysis of ΔEGF_N1, NICD constructs and empty vector. Expressed protein was detected by probing the western blot with an antibody that recognises the myc epitope tag found within all the proteins. Renilla luciferase is shown as a loading control. The position of molecular weight markers is shown in KDa. **(C)** Immunofluorescence analysis of ΔEGF_N1 and NICD proteins. Scale bar is 25 μm. **(D)** Identifying signaling activities of the Notch constructs in HEK293T cells using a Notch reporter plasmid 10×Rbpj-luc. pRL-CMV was used as a transfection control. Experiments were performed in triplicate and cells were lysed 48h post transfection. Data are presented as mean fold change (+/− SEM) in RLU (NS *P>*0.05; \*\*\**P*<0.001 one-way ANOVA and Tukey’s post-hoc test, N=3).

**Figure 2.**
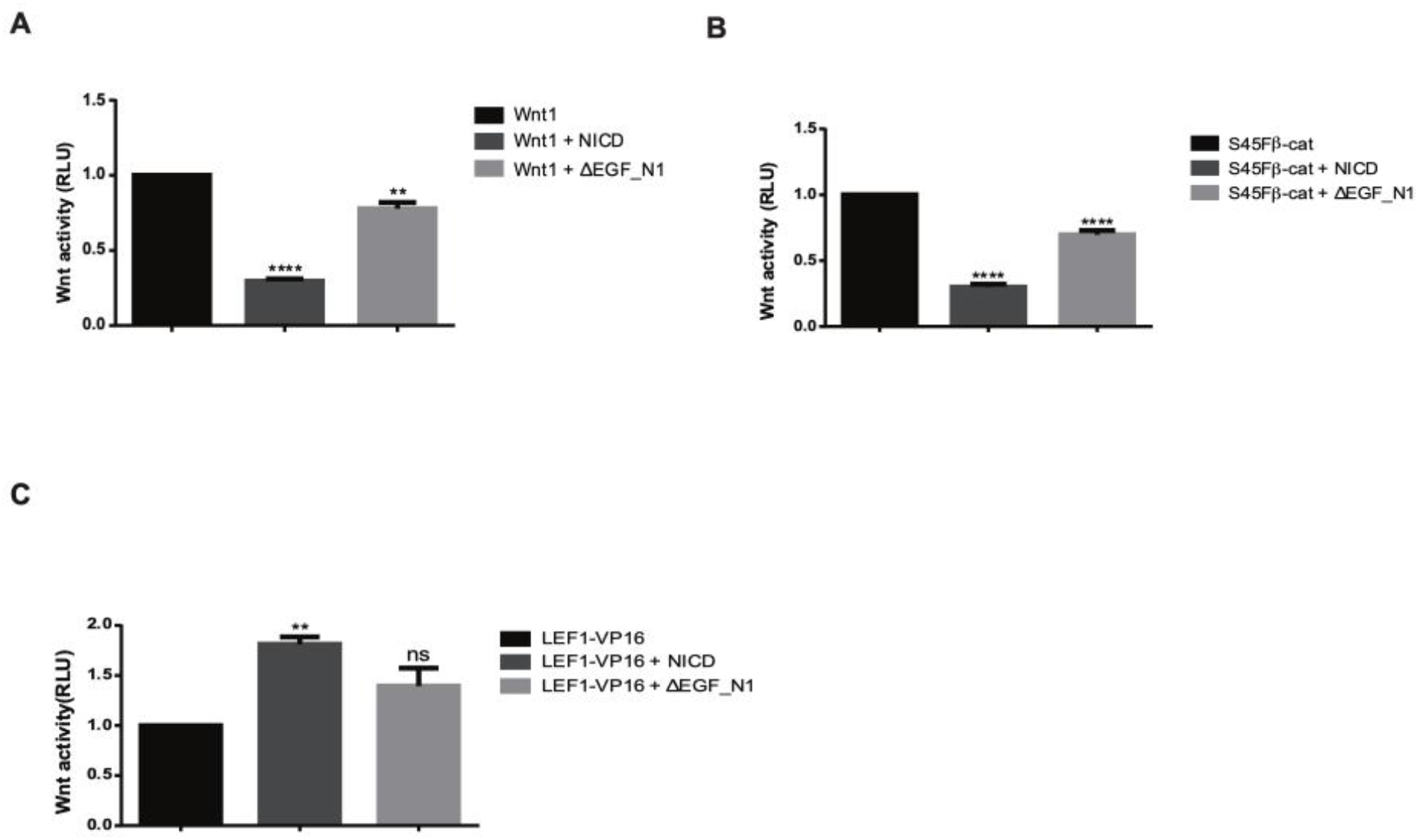
Notch signalling inhibits Wnt activity. **(A, B)** Notch inhibits Wnt1 and S45Fβ-catenin-driven Wnt signalling whereas **(C)** while neither NICD nor ΔEGF_N1 could inhibit Wnt signalling. HEK293T cells were transfected with Wnt reporter plasmid TCFAdTATA and pRL-CMV as a control **(A, B, C)**. Wnt signaling was activated by co-expression of Wnt1 **(A)** or an active form of β-catenin (S45Fβ-catenin) **(B)**, or LEF1-VP16 **(C)** alone or in the presence of ΔEGF_N1, NICD constructs as indicated. Experiments were performed in triplicate and cells were lysed 48h post transfection. Data are presented as mean fold change (+/− SEM) in RLU (NS *P>*0.05; \*\**P*<0.01, \*\*\**P*<0.001 one-way ANOVA and Tukey’s post-hoc test, N=3).

### ΔEGF_N1 requires receptor trafficking to reduce β-catenin activity

Previous work has suggested that membrane localised Notch proteins can reduce Wnt signalling by interacting with β-catenin and promoting its endocytosis and degradation (Sanders et al., 2009; Kwon et al., 2011). Consequently, we first looked to see whether the ΔEGF_N1 protein is found within vesicles of the endocytic pathway. We found clear association with Rab7 positive vesicles indicating that the protein is found within the late endosome (Fig. 3A). Additionally, expressing a dominant negative form of Vps4, which prevents trafficking from the late endosome to the lysosome (Bishop and Woodman, 2000), with ΔEGF_N1 caused a marked accumulation of the protein (Fig. 3B). This suggests that the ΔEGF_N1 is trafficked through the endocytic pathway to promote its degradation. To determine whether endocytosis was important for ΔEGF_N1 mediated suppression of Wnt signalling, we co-expressed ΔEGF_N1 with dominant negative forms of Dynamin and Rab5 and monitored changes in S45Fβ-catenin induced transcription. Dynamin and Rab5 are required for vesicle scission to separate clathrin coated pits from the plasma membrane (Damke et al., 1994) and the progression of endocytic vesicles into early endosomes (Stenmark et al., 1994) respectively, and the expression of dominant negative form of Dynamin with ΔEGF_N1 restricts the ΔEGF_N1 protein to the plasma membrane (Fig. 3C). Blocking flux through the endocytic pathway by expressing either K44A-Dynamin2 or S34N-Rab5 completely abrogated the ability of ΔEGF_N1 to reduce S45Fβ-catenin transcriptional activity (Fig. 3D&E). Interestingly, S34N-Rab5 could not attenuate the effect of NICD on S45Fβ-catenin induced transcription (Fig. S3A) indicating that the expression of these proteins is not having a non-specific effect on signalling within these cells.

**Figure 3.**
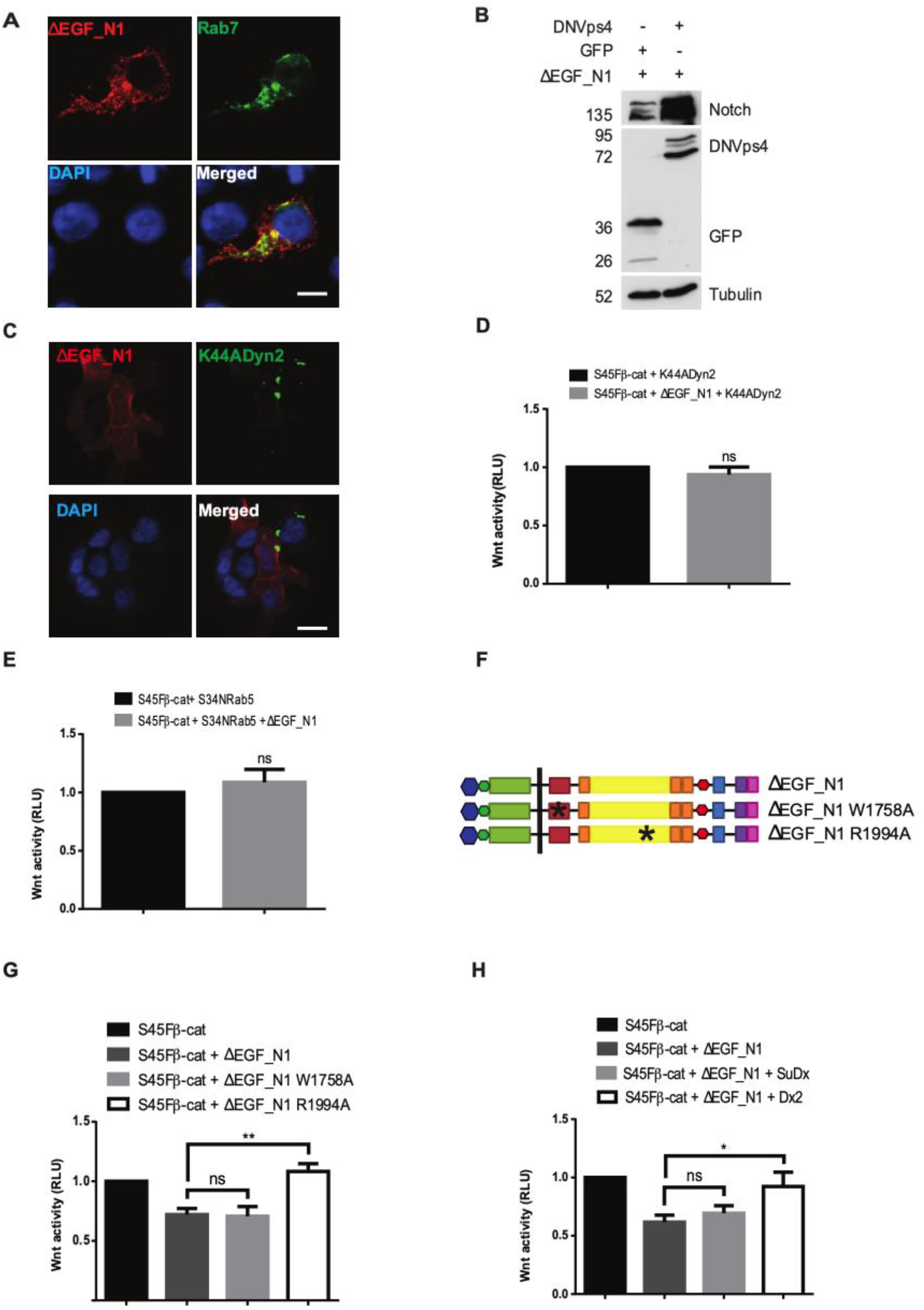
Receptor trafficking regulates ΔEGF_N1 to reduce β-catenin activity. **(A)** Membrane bound ΔEGF_N1 localises in the late endosome. Immunofluorescence analysis of ΔEGF_N1 and Rab7 proteins. Scale bar is 25 μm. **(B)** Western blot analysis of ΔEGF_N1 and DNVps4 show more ΔEGF_N1 accumulation in cells. Expressed protein was detected by probing the western blot with an antibody that recognises the myc epitope tag found within ΔEGF_N1 and GFP antibody that recognizes GFP. Tubulin is shown as a loading control. The position of molecular weight markers is shown in KDa. **(C, D, E)** Blocking endocytosis abrogates the ability of ΔEGF_N1 to inhibit S45Fβ-catenin-driven transcriptional activity. **(C)** Immunofluorescence analysis for co-localization of ΔEGF_N1 with DN form of Dynamin (K44ADyn2) protein. Scale bar is 25 μm. **(D, E)** HEK293T cells were transfected with Wnt reporter plasmid TCFAdTATA and pRL-CMV as a control. In the presence of DN form of Dynamin (K44ADyn2) **(D)** and DN form of Rab5 (S34NRab5) **(E)**ΔEGF_N1 was unable to inhibit the S45Fβ-catenin-driven transcriptional activity. **(F)** Schematic showing the structure of the ΔEGF_N1, ΔEGF_N1 W1758A and ΔEGF_N1 R1994A constructs **(G, H)** ΔEGF_N1 requires Deltex to inhibit Wnt signalling. Luciferase assay with Wnt reporter plasmid TCFAdTATA and pRL-CMV as a control indicates that ΔEGF_N1 R1994A **(G)** and Deltex **(H)** eliminates the ability of ΔEGF_N1 to inhibit the S45Fβ-catenin-driven transcriptional activity while ΔEGF_N1 W1758A or Suppressor of Deltex (SuDx) has no significant effect. Experiments were performed in triplicate and cells were lysed 48h post transfection. Data are presented as mean fold change (+/− SEM) in RLU (NS *P>*0.05; \**P*<0.05, \*\**P*<0.01, \*\*\**P*<0.001 one-way ANOVA and Tukey’s post-hoc test, N=3).

Deltex has been shown to induce the endocytosis of Notch to the late endosome (Hori et al., 2004). Once within the endosome, it is the balance between Deltex and Suppressor of Deltex that controls its localisation, with Deltex causing Notch to localise to the limiting membrane (Wilkin et al., 2008), whilst Suppressor of Deltex promotes its internalisation and trafficking to the lysosome (Wilkin et al., 2004). To establish whether Deltex plays a role in the reduction of S45Fβ-catenin induced transcription by ΔEGF_N1, we generated a point mutation (Fig. 3F), R1994A, that disrupts its interaction with Deltex. As a control, we generated a second point mutation (Fig. 3F), W1758A, that will alter the interaction of ΔEGF_N1 with RBPj and MAML. These mutations did not alter the expression or their ability to activate an RBPj-dependent reporter gene (Fig. S3B&C). However, the R1994A mutation abolished the ability of ΔEGF_N1 to effect S45Fβ-catenin induced transcription, whilst the W1758A mutation had no effect (Fig. 3G). We also over expressed either Suppressor of Deltex or Deltex with ΔEGF_N1 to alter the balance between the two proteins. Expressing Deltex also eliminated the effect of ΔEGF_N1 on S45Fβ-catenin induced transcription, whilst expressing Supressor of Deltex had little effect (Fig. 3H). Altogether these data indicate that ΔEGF_N1 requires Deltex driven endocytosis from the plasma membrane to inhibit Wnt signalling.

### NICD does not require target gene transcription to reduce β-catenin function

NICD can induce the expression of many downstream which includes members of the Hes and Hey family of transcriptional repressors (Fig. S4A). This raises the possibility that the inhibition of S45Fβ-catenin induced transcription by NICD simply reflects the fact that it is through NICD downstream genes. We tested this possibility in several different ways. Firstly, we generated a point mutation in NICD, W1758A (Fig. 4A), that alters the structure and function of the NICD/RBPj/MAML transcriptional activator and significantly reduced its transcriptional activity (Fig. 4B). As a control, we also generated the R1994A mutation in NICD (Fig. 4A) which did not alter its ability to induce the expression of an RBPj-dependent reporter gene (Fig. 4B). Neither mutation altered the expression of the NICD constructs, or their ability to localise to the nucleus (Fig. S4B&C). Both NICD constructs were able to reduce S45Fβ-catenin induced transcription (Fig. 4C). Secondly, we co-expressed NICD with an increasing concentration of a dominant negative form of Hes5. Although expression of the dominant negative form of Hes5 can clearly disrupt the transcriptional repressor function of Hes5 (Fig. 4D), it did not alter the ability of NICD to inhibit S45Fβ-catenin induced transcription (Fig. 4E). Similarly, the expression of the dominant negative form of Hey1 in increasing amounts did not alter the function of NICD to block S45Fβ-catenin induced transcription (Fig. S4D). Lastly, we expressed the NICD protein in cells where RBPj had been knockdown. This significantly reduced the ability of NICD to induce the expression of an RBPj-dependent reporter gene (Fig. 4F) and prevented the ability of NICD to inhibit S45Fβ-catenin induced transcription (Fig. 4G). Together these results indicate that NICD can inhibit Wnt signalling without inducing the expression of downstream target genes.

**Figure 4.**
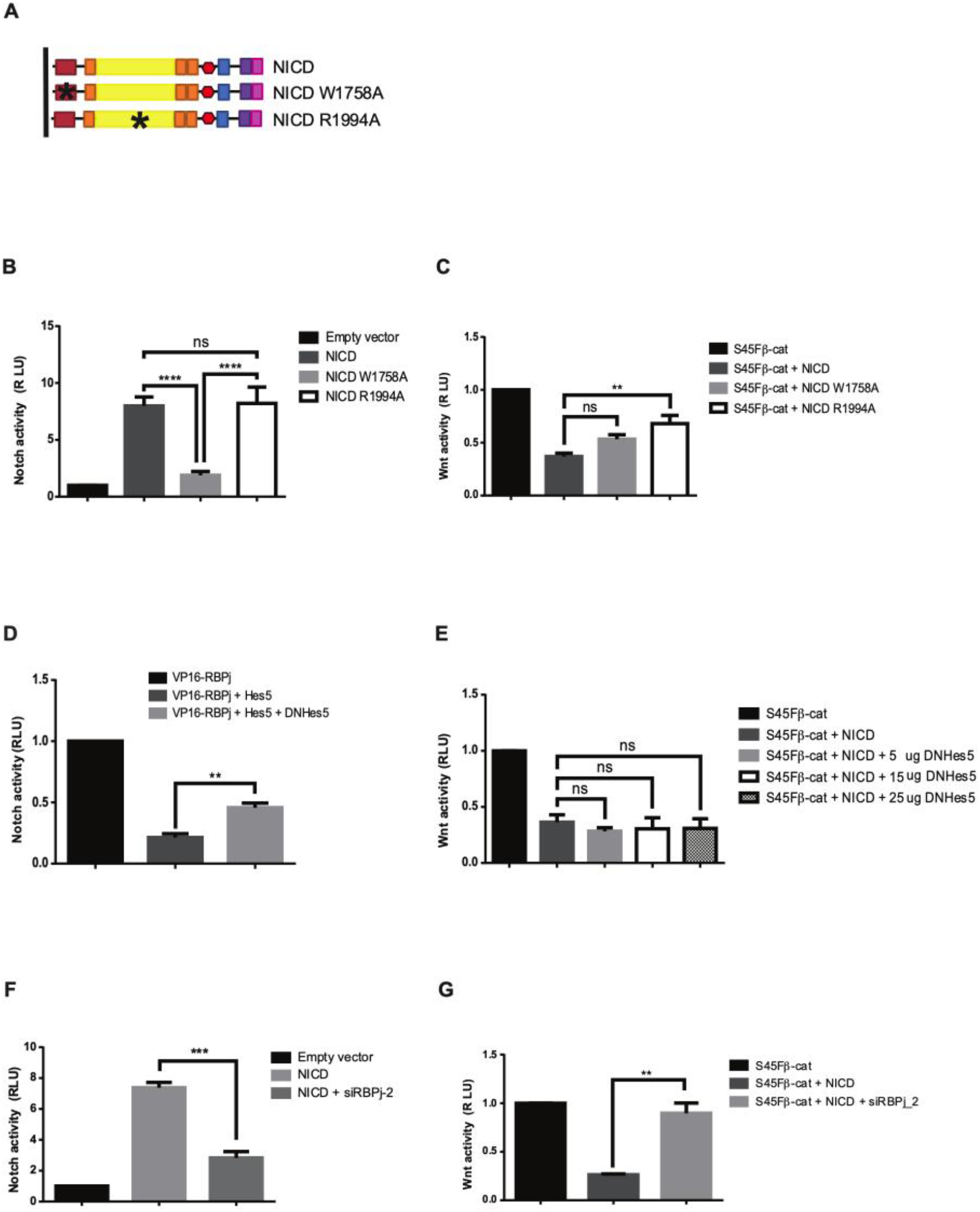
NICD reduces β-catenin function without a requirement for its target gene transcription. **(A)** Schematic showing the structure of the NICD, NICD W1758A and NICD R1994A constructs. **(B)** NICD W1758A is unable to active while NICD and NICD R1994A has no significant difference to activate the Notch signalling as assessed by luciferase assay in HEK293T cells using a Notch reporter plasmid 10xRbpj-luc. pRL-CMV was used as a transfection control. **(C)** Both of the NICD constructs reduced S45Fβ-catenin-driven transcriptional activity. Wnt reporter plasmid TCFAdTATA and a control pRL-CMV were used for the luciferase assay. **(D)** The repressor function of Hes5 is blocked by DNHes5. Notch reporter plasmid 10xRbpj-luc and a control plasmid pRL-CMV were used. **(E)** DNHes5 is not able to eliminate the ability of NICD to inhibit S45Fβ-catenin-driven transcriptional activity as assessed by Wnt reporter plasmid TCFAdTATA where the plasmid pRL-CMV was used as a control. **(F)** siRNA silencing RBPj inhibits the ability of NICD to induce Notch signaling as assessed by Notch reporter plasmid 10xRbpj-luc and a control plasmid pRL-CMV. **(G)** Inhibition of RBPj by siRNA blocked the ability of NICD to inhibit S45Fβ-catenin-driven transcriptional activity. Luciferase assay was performed using Wnt reporter plasmid TCFAdTATA and a control plasmid pRL-CMV. Experiments were performed in triplicate and cells were lysed 48h post transfection. Data are presented as mean fold change (+/− SEM) in RLU (NS *P>*0.05; \*\**P*<0.01, \*\*\**P*<0.001 one-way ANOVA and Tukey’s post-hoc test, N=3).

### Notch interacts with β-catenin to alter its function

Given the function of Hes and Hey family of proteins as transcriptional repressors, we examined the possibility of Hes5 could change the localisation of either β-catenin or members of the TCF family to prevent the formation of a transcriptional complex. To test this hypothesis, we used a split protein complementation approach (Ghosh et al., 2000), where we transfected a full length TCF4 protein fused with N-terminal half a Venus GFP (TCFV1) and a full length β-catenin protein fused with the C-terminal half of Venus GFP, (V2β catenin). Venus GFP will only refold to form a functional fluorescent protein if the TCF4 and β-catenin proteins interact (Ghosh et al., 2000). We found that in the presence of DNHes5, TCFV1 and V2β-catenin still associated to form a functional Venus GFP and DNHes5 is more potent than NICD and ΔEGF_N1 to prevent β-catenin-TCF4 complex formation and therefore Wnt signalling activation (Fig. 5A). Furthermore, the inhibitory role of Notch for Wnt signalling raised a possibility of a member of each signalling pathway to interact and prevent each other in the nucleus for their function. To test this, we used immunoprecipitation and we found a strong evidence for formation of a protein complex between NICD and S45Fβ-catenin (Fig. 5B). Moreover, both S45F and endogenous β-catenin protein levels did not change in the presence of ΔEGF_N1 (Fig. 5C). These data demonstrate that Notch forms a complex with β-catenin within the nucleus and inhibits its transcriptional activity without altering β-catenin normal expression or localisation to the nucleus.

**Figure 5.**
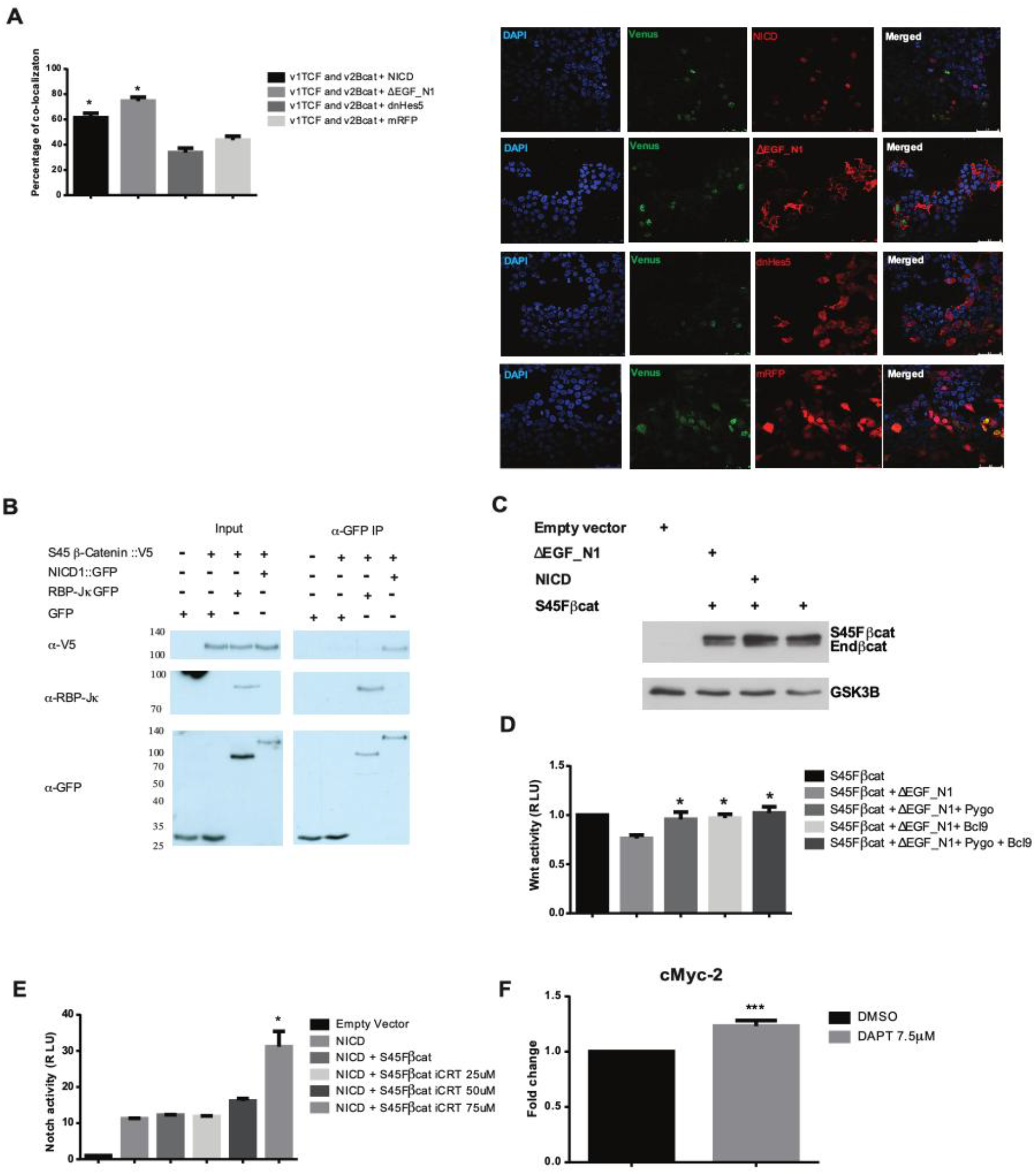
Notch and β-catenin interaction alters its function. **(A)** DNHes5 allows the formation a complex between TCFV1 and V2β-catenin. Quantification for the number of co-localization events for TCFV1 and V2β-catenin complex is measured. Immunofluorescence analysis of NICD, ΔEGF_N1, DNHes5 and RPF proteins. Scale bar is 25 μm. **(B)** A protein complex formation of β-catenin and NICD by Immunoprecipitation assay. Total nuclear lysates were used and protein detection was performed using antibodies that recognise the V5 epitope, RBPj and GFP. The position of molecular weight markers is shown in KDa. **(C)** Increased endogenous β-catenin protein levels and accumulation of S45Fβ-catenin in the cytosol. Cytosolic fraction was used to perform Western Blot analysis and antibodies that recognise endogenous β-catenin and myc-epitope were used. Glycogen synthase kinase 3 beta is shown as loading control. The position of molecular weight markers is shown in KDa. Experiments were performed in triplicate and cells were lysed 48h post transfection. Data are presented as mean fold change (+/− SEM) in RLU (NS *P>*0.05; \*\**P*<0.01, \*\*\**P*<0.001 one-way ANOVA and Tukey’s post-hoc test, N=3). **(D)** Pygo and Bcl9 prevents the ability of ΔEGF_N1 to inhibit S45Fβ-catenin-driven transcriptional activity. Luciferase assay was performed using Wnt reporter plasmid TCFAdTATA and a control plasmid pRL-CMV. **(E)** Notch signaling is enhanced when 239T cells co-expressed with NICD and S45Fβ-catenin and treated with inhibitor of β-catenin-responsive transcription (iCRT). Notch signaling activity was monitored using Notch reporter plasmid 10xRbpj-luc and a control plasmid pRL-CMV. **(F)** Quantitative PCR (qPCR) analysis of c-Myc2 gene was enhanced by DAPT treatment in HEK293T cells in comparison to DMSO treatment. Experiments were performed in triplicate and cells were lysed 48h post transfection. Data are presented as mean fold change (+/− SEM) in RLU (NS *P>*0.05; \*\**P*<0.01, \*\*\**P*<0.001 one-way ANOVA and Tukey’s post-hoc test, N=3).

We next tested the interplay between Wnt transcriptional co-activators Pygopus 1, Pygopus 2, Bcl9 and ΔEGF_N1. Inhibition of β-catenin transcriptional activity by ΔEGF_N1 was rescued in the presence of either of the Wnt transcriptional co-activators such as Pygopus or Bcl9 (Fig. 5D). This suggests the participation of Wnt downstream effectors for regulating the Wnt transcription activity when it was inhibited by the Notch. To further corroborate our findings, we used increasing amounts of β-catenin/Tcf inhibitor iCRT in the presence of forced expression of NICD and S45Fβ-catenin. Importantly, the Notch signalling activity started to increase when the iCRT concentration was increased (Fig. 5E). This finding strongly suggests the presence of ongoing interaction between Notch and Wnt downstream members and when the interaction of S45Fβ-catenin with Tcf is blocked, the freely available S45Fβ-catenin can help NICD to boost Notch signalling activity. Next, to find out whether ΔEGF-N1 was able to promote the inhibition S45Fβ-catenin-driven transcriptional activity by protein degradation, a cytosolic fractionation for S45Fβ-catenin, in the absence or presence of ΔEGF_N1, was performed. Expression of S45Fβ-catenin in HEK293T cells elevated the overall levels of endogenous β-catenin (lower band of doublet, Fig. 5C) and showed accumulation of S45Fβ-catenin (upper band of doublet, Fig. 5C) in the cytosol.

Lastly, we explored the dynamics between the endogenous expression of each of the signalling pathways by quantitative PCR analysis. We first confirmed the endogenous expression level of a known Wnt target gene cMyc-2 that was downregulated when HEK293T cells were exposed to iCRT (Fig. S5A). Importantly, when we checked the endogenous mRNA expression of c-Myc-2 when Notch signalling was inhibited and we observed that c-Myc-2 mRNA levels were upregulated in the presence of DAPT in HEK 293T cells (Fig. 5F). This indicates a role for Notch downstream effector NICD protein in mediating the transcriptional activity of Wnt pathway by regulating the expression of its endogenous gene c-Myc-2.

## Discussion

Here, we have shown that Notch inhibits Wnt signalling in two distinct ways via membrane bound ΔEGF_N1 and nuclear NICD proteins. Firstly, the interaction with Deltex mediating the sequestering of β-catenin into the membrane fraction limits the Wnt signalling activation. Secondly, NICD forms a complex with β-catenin protein within the nucleus and limits β-catenin induced transcription and Wnt signalling. Together, this study provides an evidence for the inhibitory crosstalk between Notch and Wnt pathways with in-depth mechanistic understanding for Notch protein inhibiting the Wnt signalling.

Previous studies show that Notch protein can limit Wnt signalling when membrane bound form of Notch protein is cleaved and translocated to the nucleus; however, it still remained unclear in those studies whether the initial targeting of Notch protein at the membrane was responsible for the inhibition of Wnt signalling. Our findings, in this study, provide additional evidence of Deltex-mediated trafficking causing the membrane bound Notch protein to inhibit Wnt signalling. Given the previously characterized roles of membrane associated form of Notch inhibiting Wnt signalling in Drosophila (Hayward et al., 2005; Sanders et al., 2009) and mammalian cells (Kwon et al., 2011) were independent of canonical ligand-induced Notch activation, we speculate that β-catenin may be sequestered and therefore this inhibits the Wnt signalling.

In addition to the inhibition of Wnt signalling by membrane bound Notch protein, a distinct mechanism of Wnt inhibition occurs within the nucleus by NICD transcription independent way. Consisted with previous observations (Nicolas et al., 2003; Hayward et al., 2005; Deregowski et al., 2006), NICD forms a complex with β-catenin to induce the expression of Hes and Hey; however, NICD-induced transcription activation did not alter the inhibition of Wnt signalling suggesting a role of NICD on its own or with other proteins to form a complex with β-catenin.

The dual role of Notch receptor, as an activator of the Notch signalling pathway and an inhibitor of β-catenin transcriptional activity, highlights the importance of Notch signalling pathway and its role in regulating the cell fate decisions. This has been shown by studies focusing on elucidating the a transition from a state of Notch-ON/Wnt-OFF to Notch-OFF/Wnt-ON that is often critical to progress into the differentiated state (Hayward et al., 2008; Munoz-Descalzo et al., 2012; Muñoz-Descalzo and Martinez Arias, 2012). The interplay of two states were also observed in Drosophila based on genetic perturbation experiments to demonstrate that Notch represses Wingless/Wnt during proneural cluster development. Each of these studies including ours show a requirement of a distinct separation from a Notch active or Wnt active to carefully balance the appropriate cell fate decisions.

Here, we show that Notch inhibits Wnt signalling via two distinct mechanisms. Our findings elucidate the underlying mechanisms for the Notch-ON/Wnt-OFF transition. This critical regulation holds importance to facilitate the fine-tuning activity of β-catenin and thereby the Wnt signalling. Besides a role for Wnt signalling in various developmental processes such as stem cell-renewal to lineage commitment, deregulation of the negative crosstalk and aberrant activation of either of the signalling pathways may result in disease progression. Lastly, the results presented in here may hold a promise to enhance our understanding to develop new therapeutic tools to balance signalling dynamics for robust cell fate decisions before they will be subverted in disease.

## Materials and Methods

### Cell culture

Human embryonic kidney cells that stably express the large T antigen of simian virus 40 (HEK293T) were obtained from Dr Anthony Brown (Weill Medical College, Cornell University, New York, USA) and from Dr Valerie Kouskoff (Paterson Institute for Cancer Research, Manchester, UK). HEK293T cells were cultured in DMEM medium (Lonza, Basel, Switzerland) supplemented with 10% FBS (Biowest, Nuaillé, France) and 50 μg/ml penicillin and 50 μg/ml streptomycin (Lonza, Basel, Switzerland). Cells were maintained at 37°C and 5% CO_2_ in a humidified incubator.

### Transfections and luciferase reporter assays

Cells were transfected using Lipofectamine and PLUS reagent (Invitrogen, Carlsbad, USA) or X-Treme transgene 9 transfection reagent (Roche, Basel, Switzerland), according to manufacturers’ instructions. HEK293T cells were plated at a density of 2 × 10^5^ cells/well of a 24 well plate. After 24 hr cells were transfected in triplicate with a total of 250 ng DNA cocktail containing the desired DNA plasmids and 50 ng of the Wnt reporter plasmid TCFAdTATA. As an internal control 20 ng of pRL-CMV plasmid were used. To maintain transfections with a constant amount of DNA, the plasmid pcDNA3.1(+) was used. The cell culture medium was changed 3 hr after transfection when Lipofectamine and PLUS reagent (Invitrogen, Carlsbad, USA) were used. Cells were lysed 48 hr post-transfection using 1 × Passive Lysis buffer (Promega, Madison, USA). Firefly and Renilla luciferase activities were measured using the Dual Luciferase Reporter assay system (Promega, Carlsbad, USA), according to manufacturer’s instructions, with a MicroLumatPlus plate reader (Berthold Technologies, Harpenden, UK). Data are presented as mean fold change (+/−SEM) in relative luciferase units (RLU), compared to β-catenin. Statistical analysis was perform using one-way ANOVA and Tukey’s post-hoc tests using Prism software (GraphPad, La Jolla, USA) for more than two samples or with a Student T-test for data with two samples.

### Plasmids, expression constructs and transcriptional reporters

The following plasmids were kind gifts: The **pLNCX + mWnt1** was obtained from Dr Anthony Brown (Weill Medical College, Cornell University, New York, USA). The **mβ-catenin** cDNA was obtained from Geneservice (Cambridge, UK) (I.M.A.G.E. clone 5709247) The **pcDNA3 + mN1** was obtained from Dr Jeff Nye (Northwestern University Medical School, Chicago, USA). The **pCMX + CBF1-VP16** plasmid was obtained from Dr Tasuko Honjo (Kyoto University, Japan) (Kuroda et al., 1999). The cDNA encoding mHes5 was obtained from Dr Ryoichiro Kageyama (Kyoto University, Japan). The **pCS2 + LEF-VP16** plasmid was obtained from Dr. Rolf Kemler (Max-Planck Institute for Immunology, Freiburg, Germany). The cDNA encoding **pcDNA3.1 Zeo + TCF4v1** and **pcDNA3.1 Zeo + v2β-catenin** were obtained from Dr Claudia Wellbrook (Faculty of Life Sciences, University of Manchester, UK). The **Xβ-catenin** plasmid was a gift from Dr Louise Howe (Weill Medical College of Cornell University, New York, USA). The plasmid **pcDNA3.1(+)**, used as an empty vector in transfections, was obtained from (Invitrogen, Carlsbad, USA). The **p10xCBF1-luc** Notch reporter plasmid was obtained from Dr Grahame MacKenzie (Lorantis, Cambridge, UK). The **pTOPflash** Wnt reporter was obtained from Dr Louise Howe (Weill Medical College, Cornell University, New York, USA). The **pRLCMV** reporter plasmid was obtained from (Promega, Madison, USA). The **pmRFP-C1** and **pEGFP-Rab7** plasmid were obtained from Dr Andrew Gilmore (Faculty of Life Sciences, University of Manchester, UK).

The following plasmids have already been described (Collu et al., 2012): pSecTagNC + ΔN-mN1, pSecTagNC ΔN_mN1_Δ425, pcDNA3.1 myc/HisA+ mβ-catenin.

The following plasmids where generated in our laboratory.

**pSecTagNC + ΔEGF_mN1:** the cDNA encoding mouse Notch 1 molecule that lacks all 36 EGF-like repeats was generated by cloning a PCR fragment (using mN1 4409F and mN1 4901R primers) digested HindIII/SacI along with a SacI/HindIII fragment of the mN1 cDNA into pSecTagNC+ΔEGF+LNR_mN1 (HindIII).

**pcDNA3 + mNICD:** the cDNA encoding myc tagged mNICD was generated by cloning the KpnI/BspEI fragment from pEGFP-N1 + mNICD into pcDNA3 + mN1 (KpnI/BspEI). **pcDNA3.1(+) + VP16/Rbpj:** was generated by digesting pcDNA3.1(+) KpnI/EcoRI (blunted) and ligated to remove the restriction sites from HindIII to EcoRI in the pcDNA3.1(+) multiple cloning site. The VP16-Tag was cloned from pCMX-N + VP16/Rbpj (HindIII). The Rbpj cDNA was cloned from pCMX-N + VP16/Rbpj (EcoRI).

**pcDNA3.1(+) + myc-mHes5:** was generated by digesting pcDNA3.1(+) with PmeI and inserting a fragment containing the following sequences: Kozak, myc-tag epitope, EcoRI, HindIII, and BamHI (as above), and then cloning mHes5 cDNA as an EcoRI/BamHI-digested PCR fragment generated using mHes5 73F and mHes5 663R primers.

**pmRFP + hHes5 & pmRFP + DNhHes5:** these cDNAs were generated by digesting pcDNA3.1 + myc-hHes5 and pcDNA3.1 + myc-DNhHes5 with EcoRI/BamHI and inserting the fragment into pmRFP-C1 (EcoRI/BamHI).

**pcDNA6 V5/HisA + S45Fmβ-catenin:** was generated by excising S45Fmβ-catenin from pcDNA3.1(+)/myc-HisA + S45Fmβ-catenin (generated using site directed mutagenesis, primers mβ-cat S45F F, mβ-cat S45F R) as a KpnI/XhoI fragment and inserting it into pcDNA6/V5-HisA vector (KpnI/XhoI).

**pcDNA3.1 myc/HisA + TCF4:** was generated by cloning the TCF4 cDNA from pcDNA3.1 Zeo + TCF4v1 as an EcoRI/ClaI (blunted) fragment into pcDNA3.1 myc/HisA digested with EcoRI/XhoI (blunted).

**pcDNA6 V5/HisA + TCF4:** was generated by cloning the TCF4 cDNA from pcDNA3.1 Zeo + TCF4v1 as an EcoRI/ClaI (blunted) fragment into pcDNA6 V5/HisA digested with EcoRI/XhoI (blunted).

**pGL3basic + TCFAdTATA:** this Wnt reporter plasmid was generated in two steps. Initially, p10xCBF1-luc was digested with XhoI (blunted)/BglII (blunted) and re-ligated to form pGL3basic + AdTATA. pGL3basic + AdTATA was then digested with XhoI and the SalI fragment containing the 4 TCF sites from pTOPFlash was introduced.

The following plasmids where generated by mutagenesis.

**pSecTagNC + ΔEGF_mN1 W1758A and pSecTagNC + ΔEGF_mN1 R1994A:** these plasmids were generated by PCR-based mutagenesis using the QuikChange Lightning Multi Site-Directed mutagenesis kit (Stratagene, Santa Clara, USA) using pSecTagNC + ΔEGF_mN1 as a template and primers: W1758A F and A2026VmutF, respectively.

**pcDNA3 + mNICD W1758A:** this plasmid was generated by PCR-based mutagenesis using the QuikChange Site-Directed mutagenesis kit (Stratagene, Santa Clara, USA) using pcDNA3 + mNICD as a template and the primers W1758A F and W1758A R.

**pcDNA3 + mNICD R1994A:** this plasmid was generated by PCR-based mutagenesis using the QuikChange Lightning Multi Site-Directed mutagenesis kit (Stratagene, Santa Clara, USA) using pcDNA3 + mNICD as a template and the primer A2026VmutF.

**pcDNA3.1(+) + DNhHey1, pcDNA3.1(+) + DNmHes5:** these plasmids were generated by mutating the conserved DNA binding domain Glu-Lys-X-X-Arg (EK**R) to Alanines. This generates a non-functional protein that when transfected can dimerise with endogenous proteins disrupting their function. These DN proteins were generated by PCR-based mutagenesis using the QuikChange Lightning Multi Site-Directed mutagenesis kit (Stratagene, Santa Clara, USA) using pcDNA3.1(+) + myc-hHey1, pcDNA3.1(+) + myc-mHes5 as templates and primers: hHey1 E58AK59AR62A, and mHes5 E25AK26AR29A respectively.

### S100/P100 cytosolic fractionation

Cells were washed twice in 3 ml of 1 × TBS (10 mM Tris-HCl pH 7.4 + 140 mM NaCl) + 2 mM CaCl_2_, placed on ice and lysed in 1 ml of lysis buffer (1 × TBS + freshly added protease inhibitor cocktail set I (Calbiochem, Darmstadt, Germany)) + 10 μl of 100 mM PMSF. Subsequently, cells were poured into a cold 2 ml dounce homogenizer and sheared with 30 strokes. Cell lysates were centrifuged at 1500 × g for 5 min at 4°C. The supernatant was centrifuged for 90 min at 100000 × g in a TLA 110 rotor in a Beckman Coulter Optima TLX-120 Ultracentrifuge at 4°C. After the spin, 100 μl of the supernatant, containing the cytosolic fraction, were mixed with 100 μl of 2 × Laemmli buffer (100 mM Tris-HCl pH 6.8, 200 mM DTT, 4% SDS, 0.2% bromophenol blue, 20% glycerol), boiled for 3 min and stored at −20°C.

### Western blotting

Total cell lysis and subsequent western blotting were performed as previously described (Stylianou et al., 2006). For list of the primary antibodies used see supplementary materials and methods table 1.

### Immunofluorescence

HEK293T cells were seeded on nitric acid treated coverslips at 4 × 10^5^ cells/well of a 6 well plate. Transfection was performed 24 hr later, as described above. Cells were fixed 24 hr post transfection for 10 min in 1 × PBS containing 4% formaldehyde. Following three washes with 1 × PBS, coverslips were incubated either with α-myc-Tag rabbit primary antibody (Cell signaling, Danvers, USA) for Notch constructs, diluted 1:100 in blocking solution (3% goat serum (Biosera, Sussex, UK), 0.1% Triton-X100, and 0.05% NaN_3_ in TBS) in a humidified chamber for 1 hr. Subsequently, cells were washed and coverslips were incubated with fluorescence goat α-rabbit Alexa 594 secondary antibody in blocking solution. Coverslips were mounted with VECTASHIELD mounting medium for fluorescence with DAPI H-1200 (Vector Laboratories). Images were captured with a Zeiss LSM 700, AxioObserver flexible confocal microscope (Carl Zeiss MicroImaging GmbH, Germany), using the Zeiss ZEN 2011 software (Carl Zeiss MicroImaging GmbH, Germany). The confocal software was used to determine the optimal number of Z sections when acquiring 3D optical stacks. Either maximum intensity projections of these 3D stacks or single z-section images are shown in the results. Or coverslips were mounted with Dako fluorescent mounting medium (Agilent Technologies, Santa Clara, USA). Images were collected on a Leica TCS SP5 AOBS inverted confocal using a (63× HCX PL Apo) objective. The confocal settings were as follows, pinhole 1 airy unit, scan speed 1000Hz unidirectional, format 512 × 512. Single z-section images where taken.

### Quantitative PCR

Total RNA was extracted from cells using peqGOLD TriFast solution (peqlab Biotechnologie GmbH, Erlangen, Germany) according to manufacturer’s instructions. A total of 2 μg of RNA were reverse transcribed to cDNA in a 20 μl reaction using the High Capacity RNA-to-cDNA Master Mix kit (Applied Biosystems by Life Technologies, Carlsbad, USA), following manufacturer’s instructions. The resulting cDNA was used as template for the quantitative PCR. Quantitative PCR was performed in triplicate, in a total volume of 20 μl, using the primers listed on supplementary table 4, and Fast SYBR Green PCR Master Mix (Applied Biosystems by Life Technologies, Carlsbad, USA), according to manufacturer’s instructions. The relative amounts of the PCR products were analysed using the comparative RQ method and using PPIA gene as an internal normalization control.

## Acknowledgements

We are grateful to the members of the Brennan lab for constructive discussions. We thank Dr Ahmet Ucar for critical reading manuscript and Tom Pettini for help with immunofluorescence analysis and all those who have gifted reagents. This work was supported by BBSRC and Wellcome Trust funding.

