## Supplementary Figures for "Inhibition of Wnt signalling by Notch via two distinct mechanisms"

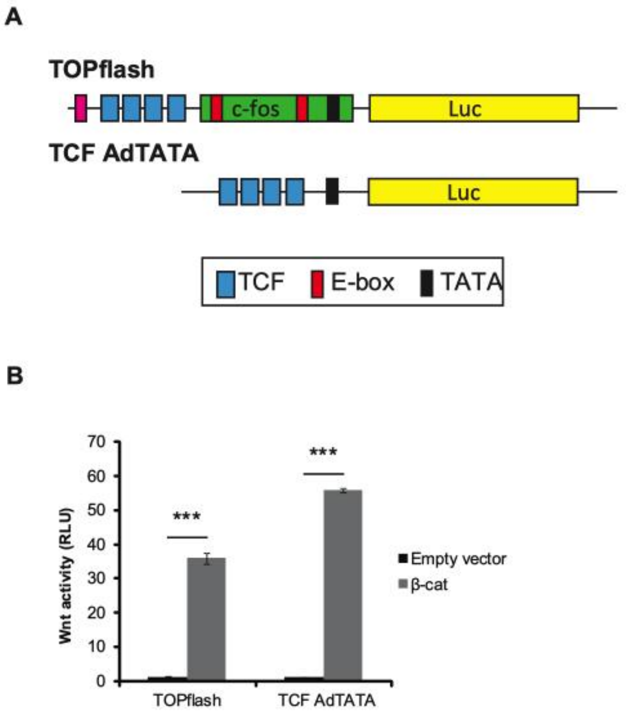


**Supplementary Figure. 1 β-catenin-induced Wnt signaling activity is greater in TCFAdTATA reporter plasmid than TOPflash reporter plasmid. (A)** Schematic showing the structure of TOPflash and TCFAdTATA reporter constructs. **(B)** Luciferase assay was performed using Wnt reporter plasmids TOPflash, TCFAdTATA and a control plasmid pRL-CMV. Experiments were performed in triplicate and cells were lysed 48h post transfection. Data are presented as mean fold change (+/- SEM) in RLU (****P*<0.001 one-way ANOVA and Tukey's post-hoc test, N=3).

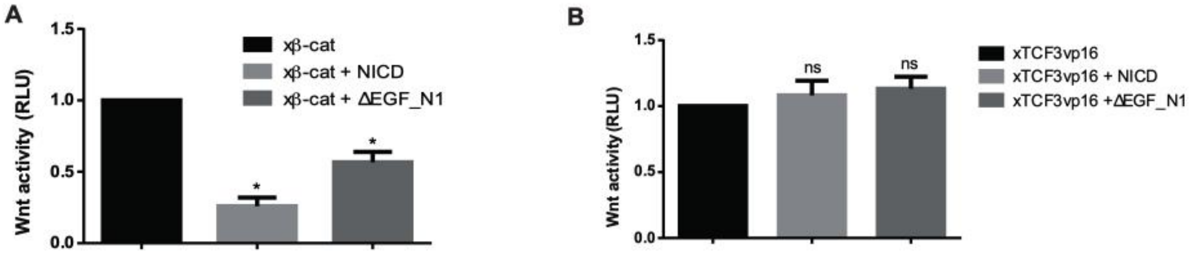


**Supplementary Fig. 2 Notch-induced Wnt signalling inhibition is conserved in Xenopus (A)** ΔEGF_N1 and NICD inhibits Xenopus β-catenin (xβ-cat)-driven transcriptional activity. **(B)** ΔEGF_N1 and NICD is unable to inhibit Xenopus TCF3VP16-driven transcriptional activity. Luciferase assays were performed using Wnt reporter plasmid TCFAdTATA and a control plasmid pRL-CMV. Experiments were performed in triplicate and cells were lysed 48h post transfection. Data are presented as mean fold change (+/- SEM) in RLU (****P*<0.001 one-way ANOVA and Tukey's post-hoc test, N=3).

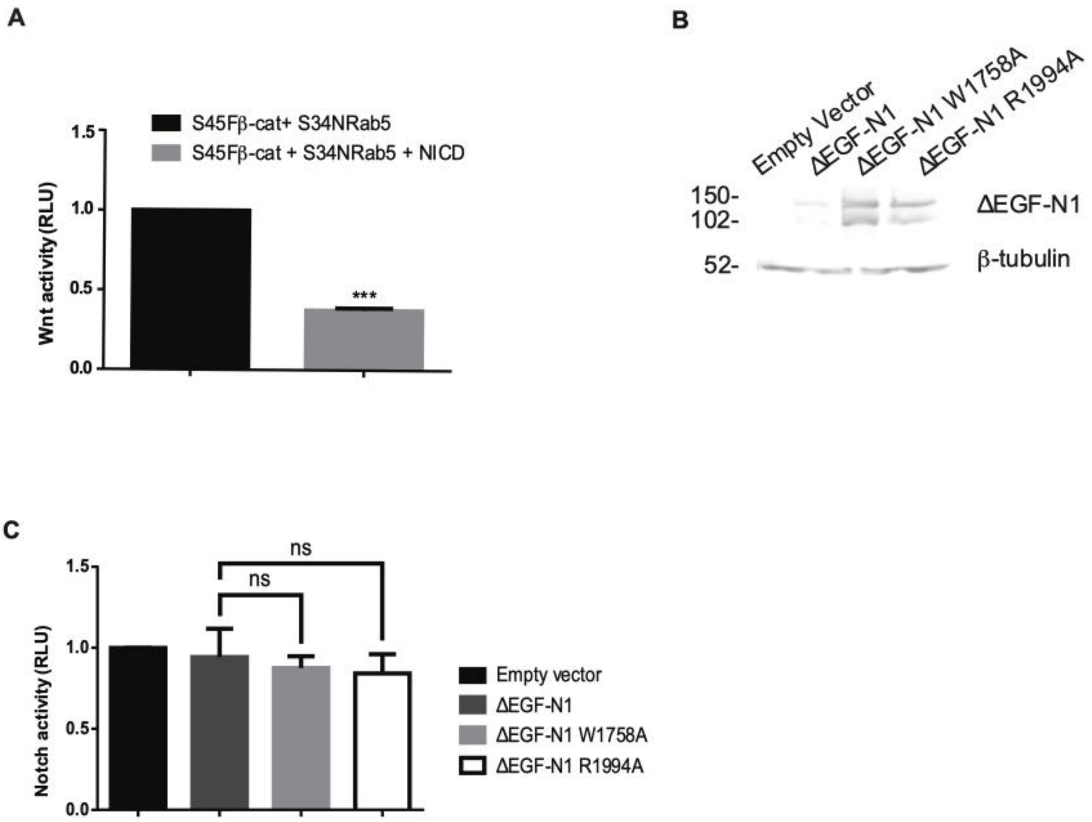


**Supplementary Figure 3 The relationship of S34NRab5 with NICD and functional characterization of ΔEGF_N1 point mutation constructs. (A)** S34NRab5 cannot attenuate the effect of NICD on S45Fβ-catenin induced transcription. Wnt reporter plasmids TCFAdTATA and a control plasmid pRL-CMV were used in luciferase assay. **(B)** The point mutations in the ΔEGF_N1 constructs did not alter their protein expression. Western blot analysis of Empty Vector, ΔEGF_N1, ΔEGF_N1, ΔEGF_N1 W1758A, ΔEGF_N1 R1994A was performed and the protein detection was achieved by probing the western blot with an antibody that recognises the myc epitope tag found within all the proteins. β-tubulin is shown as a loading control. The position of molecular weight markers is shown in KDa. **(C)** The point mutations in the ΔEGF_N1 constructs did not affect their ability to activate an RBPj-dependent Notch signalling. Luciferase assays were performed using Notch reporter plasmid 10xRbpj-luc and a control plasmid pRL-CMV. Experiments were performed in triplicate and cells were lysed 48h post transfection. Data are presented as mean fold change (+/- SEM) in RLU (NS $P > 0.05$; ***$P < 0.001$ one-way ANOVA and Tukey's post-hoc test, N=3).

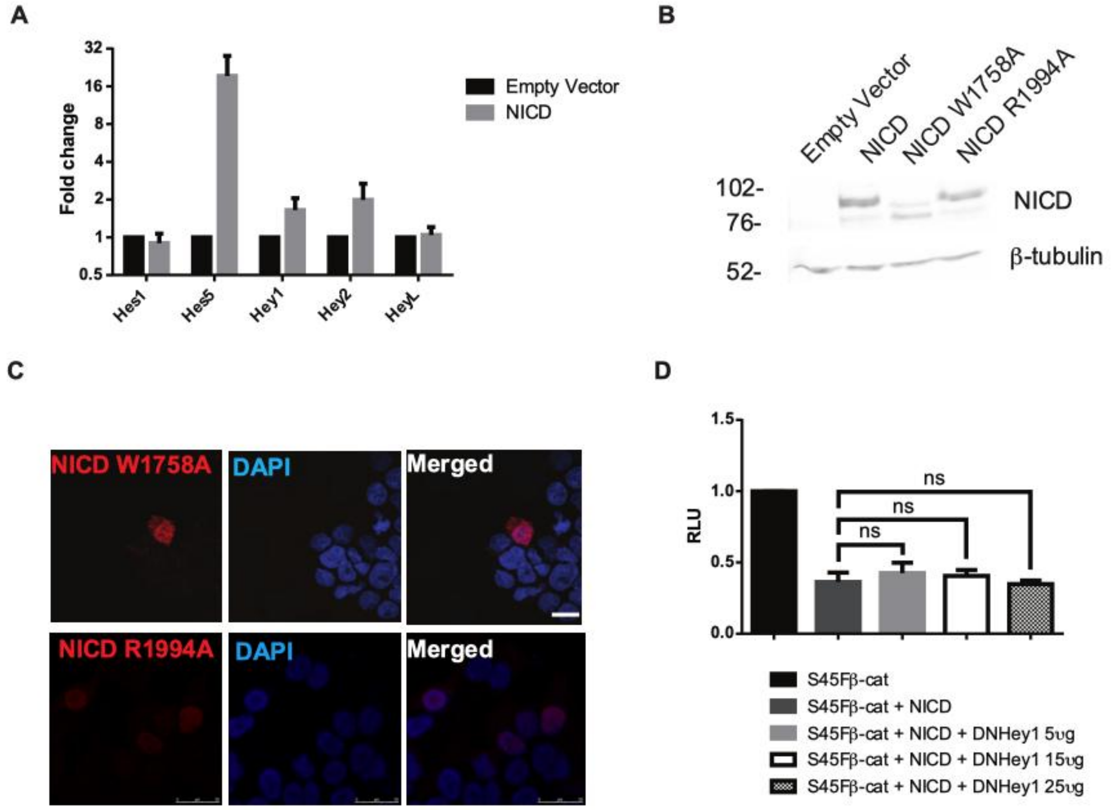


**Supplementary Figure 4 Quantitative PCR (qPCR) analysis of Notch target genes and functional characterization of NICD point mutation constructs. (A)** Amongst the Notch target genes analyzed by qPCR analysis, Hes5 is the most upregulated when NICD was overexpressed in 293T cells in comparison to induced expression of Empty Vector. **(B)** The point mutations in the NICD constructs did not alter their protein expression. Western blot analysis of Empty Vector, NICD, NICD, NICD W1758A, NICD R1994A was performed and the protein detection was achieved by probing the western blot with an antibody that recognises the myc epitope tag found within all the proteins. β-tubulin is shown as a loading control. The position of molecular weight markers is shown in KDa. **(C)** Immunofluorescence analysis of the forced exrpression of NICD W1758A and NICD R1994A proteins showed no difference in their detection. Scale bar is 25 µm. (D) DNHey1 cannot prevent the ability of NICD to inhibit S45Fβ-catenin induced transcription as assessed by the luciferase assay to monitor Wnt signalling activity using the Wnt reporter plasmid TCFAdTATA and a control plasmid pRL-CMV. Experiments were performed in triplicate and cells were lysed 48h post transfection. Data are presented as mean fold change (+/- SEM) in RLU (NS *P*>0.05; one-way ANOVA and Tukey's post-hoc test, N=3).

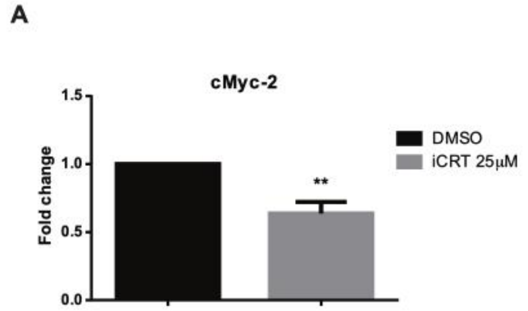


**Supplementary Fig. 5 Quantitative PCR (qPCR) analysis of cMyc-2 (A)** Quantitative PCR (qPCR) analysis of c-Myc2 gene revealed that its mRNA expression was downregulated whe HEK293T cells were treated with iCRT in comparison to DMSO treatment. Data are presented as mean fold change (+/- SEM) in RLU (***P*<0.01 one-way ANOVA and Tukey's post-hoc test, N=3).

### Supplementary information

**Table S1. Antibodies used for western blot and immunofluorescence.**

| **PRIMARY ANTIBODIES WB** | | | | |
|---|---|---|---|---|
| **Host species** | **Immunogen** | **Dilution** | **Supplier** | **Cat. Number** |
| Mouse | α-Tubulin | 1:1000 | Gift K.Gull Uni of Mcr | N/A |
| Mouse | β-catenin | 1:2000 | BD Transduction lab | 610154 |
| | GFP | 1:1000 | | |
| | HA | | | |
| Mouse | Myc (clone 4A6) | 1:1000 | Upstate | 05-724 |
| Mouse | Renilla luciferase | 1:500 | Cell Signaling | 2272 |
| Rabbit | RFP | 1:1000 | MBI | PM005 |
| Mouse | V5 | 1:1000 | Invitrogen | R960-25 |
| Mouse | VP16 | 1:200 | Santa Cruz | sc-7545 |
| **PRIMARY ANTIBODIES IF** | | | | |
| Rabbit | Myc-Tag | 1:400 | Cell signaling | 2272 |

| **SECONDARY ANTIBODIES WB** | | | | |
|---|---|---|---|---|
| **Host species** | **Immunogen** | **Dilution** | **Supplier** | **Cat. Number** |
| Donkey | Mouse IgG | 1:10000 | Jackson | 715-035-150 |
| Donkey | Rabbit IgG | 1:10000 | Jackson | 715-035-152 |
| **SECONDARY ANTIBODY IF** | | | | |
| Goat | Rabbit Alexa 594 | 1:400 | Molecular Probes | A11037 |

**Table S2. PCR and sequencing primers**

| **Primer name** | **Sequence (5'-3')** |
|---|---|
| mN1 4409F | GGTAAAGCTTCAGATTGAGGAGGCATGTGAG |
| mN1 4901R | GCTTGAAGACCACGTTGGTGT |
| mN1 5042F | TAGTAAGCTTGAGCTGGACCCTATGGACAT |
| mN1 5589R | TGCTGCTGAGTCCACTGTCT |

| | |
|---|---|
| mHes5 73F | TAGCGAATTCTGGCATGGCACCTAGTACCGTGG |
| mHes5 663R | TCGTGGATCCTGAACTGCGGCTGGGGAATGTC |
| hHey1 98 F | TAGCGAATTCTATGAAGCGTGCTCACCCCGAGT |
| hHey1 1077 R | TAGCGGATCCTCTTAGCAACAGTCCAGCCCA |

**Table S3. Mutagenesis primers.**

| Primer name | Sequence (5’-3’) |
|---|---|
| mβ-cat S45FF | ACCACCACAGCTCCTTTCCTGAGTGGCAAGGGC |
| mβ-cat S45FR | GCCGTTGCCACTCAGGAAAGGAGCTGTGGTGGT |
| W1758A F | CAGCATGGCCAGCTC**GCG**TTGCCTGAGGGTTTG |
| W1758A R | GAAACCCTCAGGGAA**CGC**GAGCTGGCCATGCTG |
| R1994AmutF | TTGCATTGGGCGGCC**GTG**GTGAACAATGTGGAT |
| hHey1E58AK59AR62A | GGAGAGGAATAATT**GCGGCG**CGCCGA**GCA**GACCGGATCA ATAAC |
| mHes5E25AK26AR29A | GGAAGCCGGTGGTG**GCGGCG**ATGCGT**GCG**GACCGCATCA ACAGC |

**Table S4. Quantitative PCR primers.**

| Primer name | Sequence (5’-3’) |
|---|---|
| hPygopus1-a F | GTTTCCTCGCATGGTGGTGA |
| hPygopus1-a R | TGGATTCGGTGGTGGAGCAT |
| hPygopus2-a F | GCAAGGCCGGTCTGCAAATG |
| hPygopus2-a R | GGGTGGTGCAAACTCCGTCA |
| hBcl9-a F | TGTGGCCAGCTCAGATGACG |
| hBcl9-a R | AACCACGGGGTTTGGACCTG |
| hHey1 1-449 F | AAAGCGTGAGCGGGATCAG |
| hHey1 1-449 R | CTCGTCGGCGCTTCTCAAT |
| hHey1-real5' | GGCAGGAGGGAAAGGTTACT |
| hHey1-real3' | GCTGGGAAGCGTAGTTGTTG |
| hHey2-real5' | AAGATGCTTCAGGCAACAGG |
| hHes1-real5' | TCTGAGCCAGCTGAAAACAC |

| | |
|---|---|
| hHes1-real3' | CTCGGTACTTCCCCAGCAC |
| hHes5-real5'-new | CCCAAAGAGAAAAACCGACTG |
| hHes5-real3'-new | GCTTGGAGTTGGGCTGGT |
| hHeyL-real5' | AGACCGCATCAACAGTAGCC |
| hHeyL-real3' | CAAAGAATCCTGTCCCACCA |
| hPPIA-real5' | ATGCTGGACCCAACACAAA |
| hPPIA-real3' | TTTCACTTTGCCAAACACCA |
